# Comparative Genome Analysis of ‘*Candidatus* Phytoplasma luffae’ Reveals the Influential Roles of Potential Mobile Units in Phytoplasma Evolution

**DOI:** 10.1101/2021.09.09.459700

**Authors:** Ching-Ting Huang, Shu-Ting Cho, Yu-Chen Lin, Choon-Meng Tan, Yi-Ching Chiu, Jun-Yi Yang, Chih-Horng Kuo

**Affiliations:** Institute of Plant and Microbial Biology, Academia Sinica, Taipei 115, Taiwan; Graduate Institute of Biochemistry, National Chung Hsing University, Taichung 402, Taiwan; PhD Program in Microbial Genomics, National Chung Hsing University and Academia Sinica, Taichung 402, Taiwan; Advanced Plant Biotechnology Center, National Chung Hsing University, Taichung 402, Taiwan; Molecular and Biological Agricultural Sciences Program, Taiwan International Graduate Program, National Chung-Hsing University and Academia Sinica, Taipei 115, Taiwan; Biotechnology Center, National Chung Hsing University, Taichung 402, Taiwan

**Keywords:** Plant pathogen, phytoplasma, genomics, molecular evolution, mobile genetic element, effector

## Abstract

Phytoplasmas are insect-transmitted plant pathogens that cause substantial losses in agriculture. In addition to economic impact, phytoplasmas induce distinct disease symptoms in infected plants, thus attracting attention for research on molecular plant-microbe interactions and plant developmental processes. Due to the difficulty of establishing an axenic culture of these bacteria, culture-independent genome characterization is a crucial tool for phytoplasma research. However, phytoplasma genomes have strong nucleotide composition biases and are repetitive, which make it challenging to produce complete assemblies. In this study, we utilized Illumina and Oxford Nanopore sequencing technologies to obtain the complete genome sequence of ‘*Candidatus* Phytoplasma luffae’ strain NCHU2019 that is associated with witches’ broom disease of loofah (*Luffa aegyptiaca*) in Taiwan. The fully assembled circular chromosome is 769 kb in size and is the first representative genome sequence of group 16SrVIII phytoplasmas. Comparative analysis with other phytoplasmas revealed that NCHU2019 has an exceptionally repetitive genome, possessing a pair of 75 kb repeats and at least 13 potential mobile units (PMUs) that account for ~25% of its chromosome. This level of genome repetitiveness is exceptional for bacteria, particularly among obligate pathogens with reduced genomes. Our genus-level analysis of PMUs demonstrated that these phytoplasma-specific mobile genetic elements can be classified into three major types that differ in gene organization and phylogenetic distribution. Notably, PMU abundance explains nearly 80% of the variance in phytoplasma genome sizes, a finding that provides a quantitative estimate for the importance of PMUs in phytoplasma genome variability. Finally, our investigation found that in addition to horizontal gene transfer, PMUs also contribute to intra-genomic duplications of effector genes, which may provide redundancy for neofunctionalization or subfunctionalization. Taken together, this work improves the taxon sampling for phytoplasma genome research and provides novel information regarding the roles of mobile genetic elements in phytoplasma evolution.

## Introduction

Phytoplasmas are uncultivated bacteria associated with plant diseases in several hundred species (Lee et al., 2000; Hogenhout et al., 2008; Bertaccini et al., 2014; Namba, 2019). In infected plants, phytoplasma cells are restricted to phloem tissues and can secrete effector proteins that cause developmental abnormalities of the hosts (Sugio et al., 2011b). Typical symptoms of phytoplasma infections include stunting, dwarfism, virescence (i.e., greening of flowers), phyllody (i.e., abnormal development of floral parts into leaf-like tissues), and witches’ broom (i.e., proliferation of stems and leaves), which result in substantial agricultural losses.

For classification of these uncultivated bacteria, a system based on restriction fragment length polymorphism (RFLP) analysis of their 16S rRNA genes was developed in the 1990s (Lee et al., 1993, 1998; Gundersen and Lee, 1996) and at least 33 16S rRNA gene RFLP (16Sr) groups have been described (Zhao et al., 2009; Zhao and Davis, 2016). Later, a provisional genus-level taxon ‘*Candidatus* Phytoplasma’ was proposed to accommodate these bacteria (The IRPCM Phytoplasma/Spiroplasma Working Team - Phytoplasma taxonomy group, 2004) and at least 41 ‘*Ca*. P.’ species have been described or proposed (Bertaccini and Lee, 2018). Based on analysis of 16S rRNA and other conserved genes, phytoplasmas are divided into three major phylogenetic clusters (Hogenhout and Seruga Music, 2009; Chung et al., 2013; Seruga Music et al., 2019). Early genomics studies were mainly conducted for clusters I (i.e., ‘*Ca*. P. asteris’ of group 16SrI and ‘*Ca*. P. australiense’ of 16SrXII) (Oshima et al., 2004; Bai et al., 2006; Tran-Nguyen et al., 2008) and II (i.e., ‘*Ca*. P. mali’ of 16SrX) (Kube et al., 2008). In comparison, cluster III harbors the highest level of diversity, yet has received limited attention for comparative genomics studies (Chung et al., 2013; Wang et al., 2018a).

To improve our understanding of phytoplasma genome diversity, we conducted whole genome sequencing of a ‘*Ca*. P. luffae’ strain collected in Taiwan. The species-level taxon ‘*Ca*. P luffae’ belongs to group 16SrVIII in cluster III and is associated with witches’ broom disease of loofah (*Luffa aegyptiaca*) (Davis et al., 2017). The availability of a complete genome sequence from this taxon provides a complete view of its gene content, which can facilitate the study of its pathogenesis mechanisms and other aspects of its biology. More importantly, with the increased availability of genome sequences from diverse phytoplasmas (**Table 1**), we performed genus-level comparative analysis to obtain a more comprehensive picture of their genomic diversity. This improves upon previous works that are limited to comparisons of closely related taxa or have sparse sampling (Bai et al., 2006; Tran-Nguyen et al., 2008; Kube et al., 2012; Saccardo et al., 2012; Andersen et al., 2013; Chung et al., 2013; Orlovskis et al., 2017; Wang et al., 2018a; Cho et al., 2019, 2020a; Seruga Music et al., 2019; Kirdat et al., 2021; Zhao et al., 2021). Furthermore, our focused analysis of the potential mobile units (PMUs) (Bai et al., 2006) revealed the influential roles of these mobile genetic elements in the evolution of phytoplasma genome organization and effector gene content.

**Table 1.**
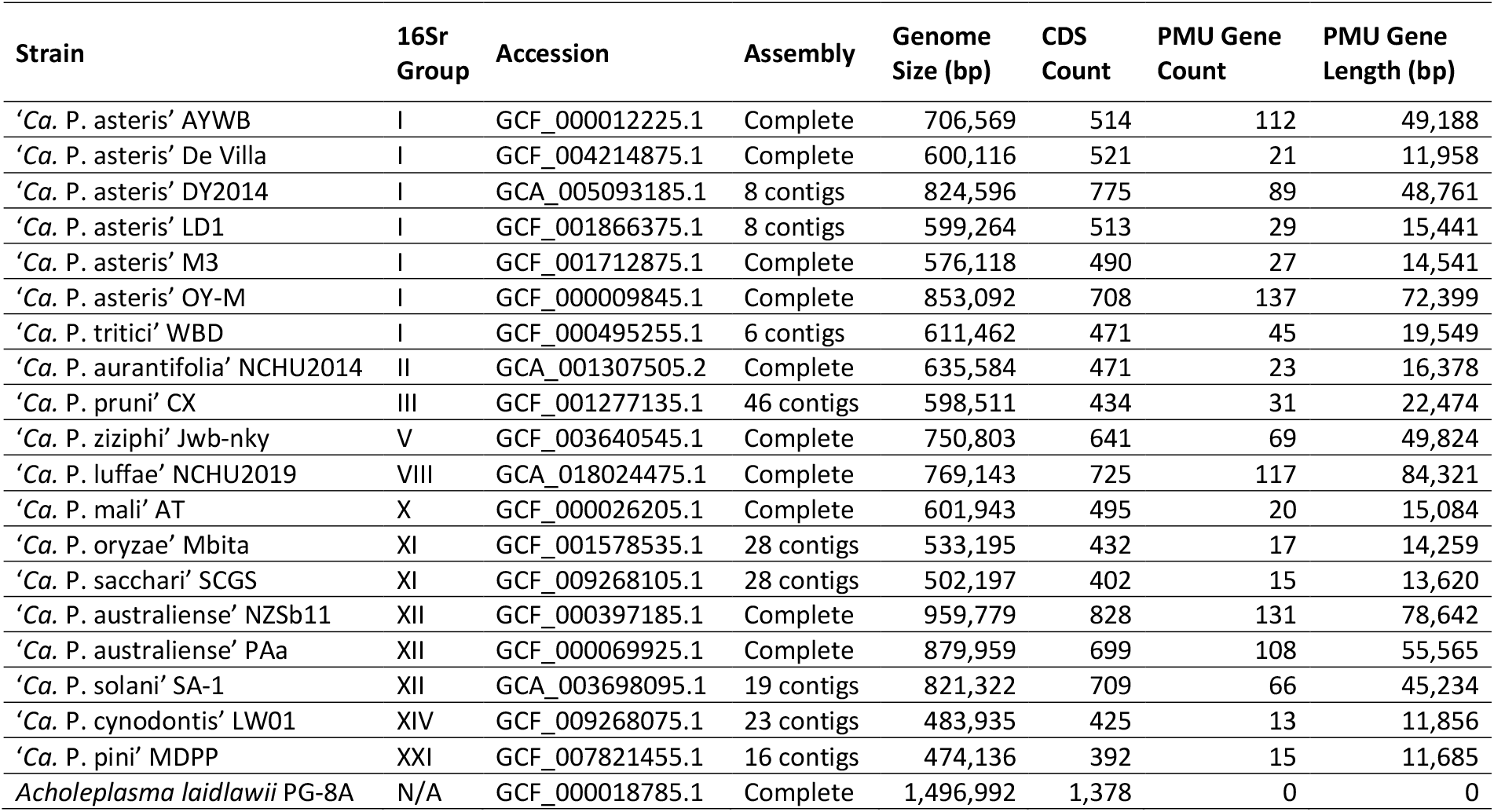
List of the genome assemblies analyzed. For each strain, information regarding the 16S rRNA gene (16Sr) group, genome accession number, assembly status, genome size, coding sequence (CDS) count, potential mobile unit (PMU) gene count, and combined length of PMU genes are provided. The PMU gene information is based on homologs of eight core genes (i.e., *tra5*, *dnaB*, *dnaG*, *tmk*, *hflB*, *himA*, *ssb*, and *rpoD*); other genes such as those encode hypothetical proteins or putative secreted proteins are excluded. *Acholeplasma laidlawii* is included as an outgroup.

## Materials and Methods

### Biological Materials

The strain NCHU2019 was collected from a naturally infected loofah plant found in Dacheng Township (Changhua County, Taiwan; 23.861860 N, 120.291322 E) on July 4th, 2019 (**Figure 1A**). After initial collection, the bacterium was transferred to lab-grown loofah plants (cultivar A-Jun, Known-You Seed Co., Kaohsiung, Taiwan) through grafting and maintained in a plant growth facility in the National Chung Hsing University (Taichung, Taiwan) (**Figure 1B**). To confirm the presence and identity of this phytoplasma strain, a partial sequence of the rRNA operon was PCR amplified using the phytoplasma-specific primer set P1/P7 for Sanger sequencing as described (Chung et al., 2013).

**Figure 1.**
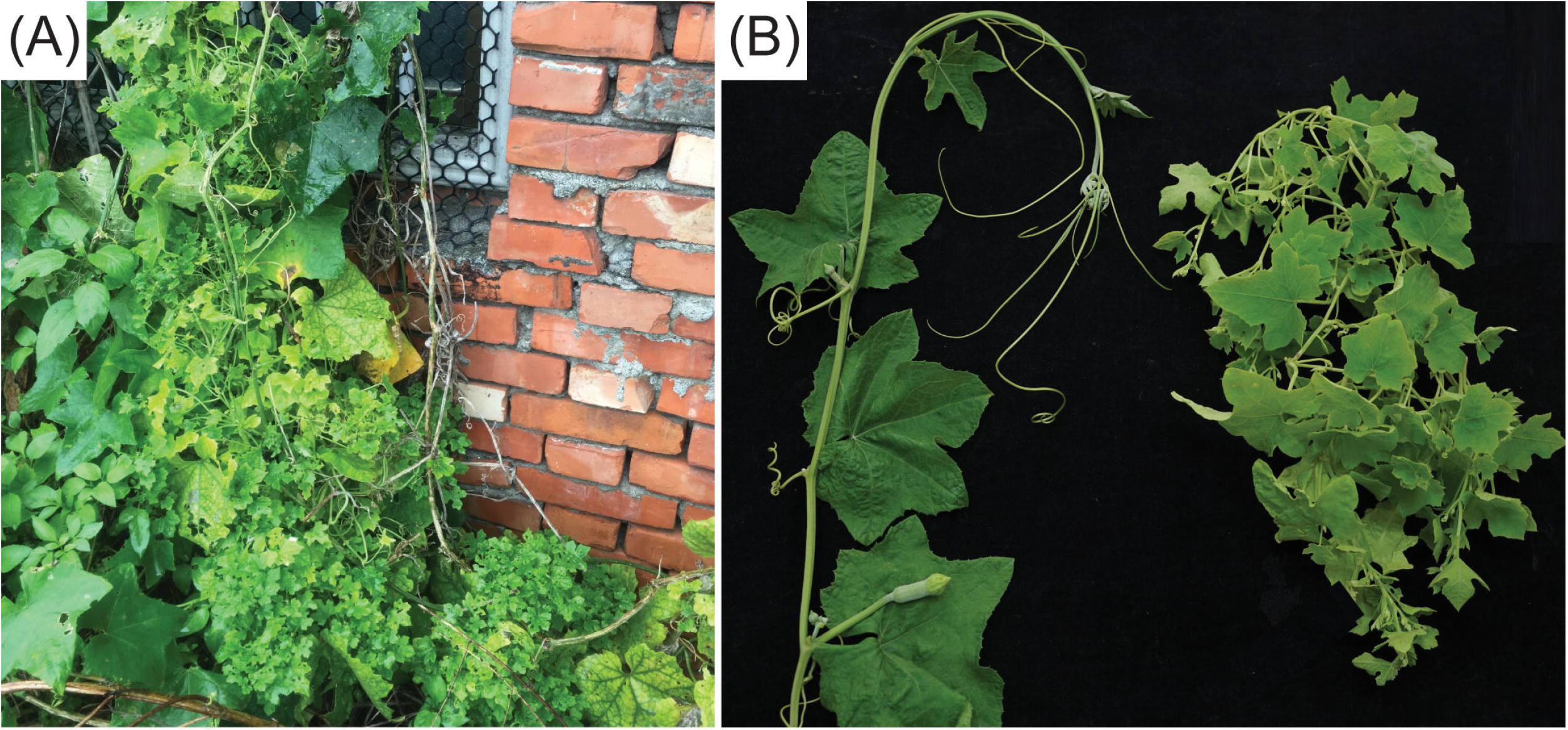
Infection symptoms. (A) The loofah plant exhibiting phytoplasma infection symptoms that was collected in Changhua, Taiwan. (B) Loofah plants grown in the lab. Left, healthy control; right, grafted with the infected plant shown in panel A.

### Genome Sequencing, Assembly, and Annotation

The procedures for genome sequencing and analysis were based on those described in our previous work on phytoplasma genomes (Chung et al., 2013; Cho et al., 2019, 2020a, 2020b). All kits were used according to the manufacturer’s protocols and all bioinformatics tools were used with the default settings unless stated otherwise.

For whole genome shotgun sequencing, leaves from one artificially infected plant exhibiting typical symptoms (i.e., small leaves and witches’ broom) (**Figure 1B**) were collected for total genomic DNA extraction using the Wizard Genomic DNA Purification Kit (A1120; Promega, USA). For Illumina sequencing, the DNA sample was processed using the KAPA Library Preparation Kit (KK8234) and the Invitrogen SizeSelect Gels (G6610-02) to obtain ^~^550 bp fragments, followed by MiSeq 2×300 bp paired-end sequencing (v3 chemistry). For Oxford Nanopore Technologies (ONT) sequencing, the library was prepared using the ONT Ligation Kit (SQK-LSK109) without shearing or size selection, followed by MinION sequencing (R9.4.1 chemistry) and Guppy v3.3.3 base-calling.

The *de novo* genome assembly involved two stages. In the first stage, only the Illumina reads were used for running Velvet v1.2.10 (Zerbino and Birney, 2008) with k-mer length set to 151 and minimum contig length set to 2,000 bp. To identify putative phytoplasma contigs, all contigs were was queried against the NCBI non-redundant protein database (Benson et al., 2018) using BLASTX v2.10.0 (Camacho et al., 2009). Those have a hit with an e-value of < 1e-15 to phytoplasma proteins were selected for manual inspection. False positive contigs derived from plant chloroplast and mitochondrial genomes were identified based on the difference in sequencing depth and verified based on BLASTN v2.10.0 (Camacho et al., 2009) searches against the NCBI standard nucleotide collection database (Benson et al., 2018).

To start the second stage of assembly, the putative phytoplasma contigs identified from the first stage were used as the reference for extracting phytoplasma reads from the Illumina data set using BWA v0.7.17 (Li and Durbin, 2009) with an alignment score cutoff of 30 and from the ONT data set using Minimap2 v2.15 (Li, 2018) with an alignment score cutoff of 1,000. The extracted reads were processed together for *de novo* hybrid assembly using Unicycler v0.4.9b. The contigs were validated as belonging to the phytoplasma genome by BLAST searches according to the process utilized in the first stage and used as the starting point for an iterative process of assembly improvement. In each iteration, all Illumina and ONT raw reads were mapped to the draft assembly as described. The mapping results were programmatically checked using the “mpileup” function of SAMTOOLS v1.9 (Li et al., 2009) and manually inspected using IGV v2.5.0 (Robinson et al., 2011) to identify possible assembly errors. Regions with raw reads mapping results exhibiting abnormalities were cut and re-arranged manually based on the continuity of ONT long reads, then validated using the mapping results in the next iteration. During the early iterations, the mapping results of ONT reads were used to provide scaffolding information and validate the overall organization of the circular chromosome, particularly the junctions between repetitive and unique regions. Additionally, the reads mapped to contig ends were visually inspected using IGV for manual selection of representative reads that can extend contigs and fill gaps. During the later iterations, the mapping results of Illumina reads were used to validate bp-scale indels and possible sequencing errors introduced by the ONT reads. The process was repeated until the complete genome assembly was obtained and all regions are supported by the raw reads mapping results. Additionally, the “depth” function of SAMTOOLS is used to calculate the sequencing coverage.

To provide a genome size estimate based on k-mer distribution, all Illumina reads mapped to the finalized assembly with an alignment score above 200 were extracted. Based on these reads, occurrences of k-mers in the size range between 17 and 63 were calculated by using jellyfish v2.2.8 (Marçais and Kingsford, 2011). The genome size was estimated by dividing the total k-mer count with the peak depth as suggested previously (Lu et al., 2016).

The procedure of gene prediction was performed using RNAmmer v1.2 (Lagesen et al., 2007), tRNAscan-SE v1.3.1 (Lowe and Eddy, 1997), and Prodigal v2.6.3 (Hyatt et al., 2010). The annotation was based on the homologous gene clusters present in other phytoplasma genomes (**Table 1**) as identified by BLASTP v 2.10.0 (Camacho et al., 2009) and OrthoMCL v1.3 (Li et al., 2003), followed by manual curation based on information obtained from GenBank (Benson et al., 2018), KEGG (Kanehisa et al., 2010), and COG (Tatusov et al., 2003) databases. Additionally, putative secreted proteins were predicted using SignalP v5.0 (Armenteros et al., 2019) based on the Gram-positive bacteria model. Those candidates with transmembrane domains as identified by TMHMM v2.0 (Krogh et al., 2001) were removed and the remaining ones were required to have a signal peptide length in the range of 21 to 52 amino acids. For visualization, the Circos v0.69-6 (Krzywinski et al., 2009) was used to draw the genome map.

### Comparative Analysis

For comparative analysis with other representative phytoplasma genomes (**Table 1**), homologous gene clusters were identified using OrthoMCL (Li et al., 2003). Multiple sequence alignments of homologous genes were prepared using MUSCLE v3.8.31 (Edgar, 2004) and visualized using JalView v2.11 (Waterhouse et al., 2009). Maximum likelihood phylogenies were inferred using PhyML v3.3 (Guindon and Gascuel, 2003) with the default LG substitution model and visualized using FigTree v1.4.4. PHYLIP v3.697 (Felsenstein, 1989) was used for bootstrap analysis. Classification of the 16S RFLP group was performed using *i*PhyClassifier (Zhao et al., 2009).

For whole-genome comparison, FastANI v1.1 (Jain et al., 2018) was used to calculate the proportion of genomic segments mapped and the average nucleotide identity (ANI). For pairwise genome alignments, Mummer v3.23 (Kurtz et al., 2004) was used with the options “--maxmatch --mincluster 30” and the results were visualized using genoPlotR v0.8.9 (Guy et al., 2010).

The PMU analysis was based on the eight core genes (i.e., *tra5*, *dnaB*, *dnaG*, *tmk*, *hflB*, *himA*, *ssb*, and *rpoD*) defined previously (Bai et al., 2006). To ensure uniform annotation and to include possible pseudogenes, all genome assemblies were examined by using available PMU genes sequences as queries to run TBLASTN searches. All statistical tests were performed in the R statistical environment (R Core Team, 2019); correlation coefficients were calculated using the ‘cor.test’ function, linear regression was performed using the ‘lm’ function and visualized using the ‘plot’ and ‘abline’ functions.

## Results and Discussion

### Genomic Characterization of NCHU2019

The shotgun sequencing generated ^~^4.7 Gb of Illumina raw reads and ^~^3.5 Gb of ONT raw reads. In the first stage of *de novo* assembly with only the Illumina reads, a draft assembly with 49 contigs totaling 450,754 bp was obtained (longest contig: 31,507 bp; shortest contig: 2,001 bp; N50: 12,506 bp). Based on the mapping results to this first draft assembly, we extracted 0.9% (136,116 out of 15,655,220) of all Illumina raw reads and 1.7% (9,416 out of 539,354) of all ONT raw reads. These reads correspond to a sequencing depth of 71- and 110-fold, respectively. In the second stage of *de novo* hybrid assembly using these extracted reads from both sequencing platforms, a second draft assembly with 10 contigs totaling 668,430 bp was obtained. These 10 contigs include two long ones that are 552,199 and 103,217 bp in size, as well as eight short ones that are shorter than 3,000 bp. Examination of the mapping results to this second draft assembly found 186,979 Illumina reads with 73-fold coverage and 13,242 ONT reads with 95-fold coverage.

During the iterative assembly improvement process, a pair of duplicated chromosomal segments that are each ^~^75 kb in size and multiple smaller repeats were identified. These regions were manually corrected to generate the finalized assembly. All junctions involving repetitive regions were verified based on visual inspection of the ONT long read mapping results. For the finalized assembly, we obtained a circular contig that corresponds to the phytoplasma chromosome; no plasmid was found. This circular contig is 769,143 bp in size with 23.3% G+C content (**Figure 2**). For the final round of raw reads mapping examination, 149,857 Illumina reads and 13,649 ONT reads were mapped to this circular contig, corresponding to 51- and 80-fold sequencing depth, respectively. The decrease of sequencing depth compared to the draft assemblies is likely explained by the repetitive regions being resolved. As a support for this inference, the sequencing depth exhibits a nearly uniform distribution across the entire finalized circular contig, with repetitive regions having similar depth as non-repetitive regions (**Figure S1**).

**Figure 2.**
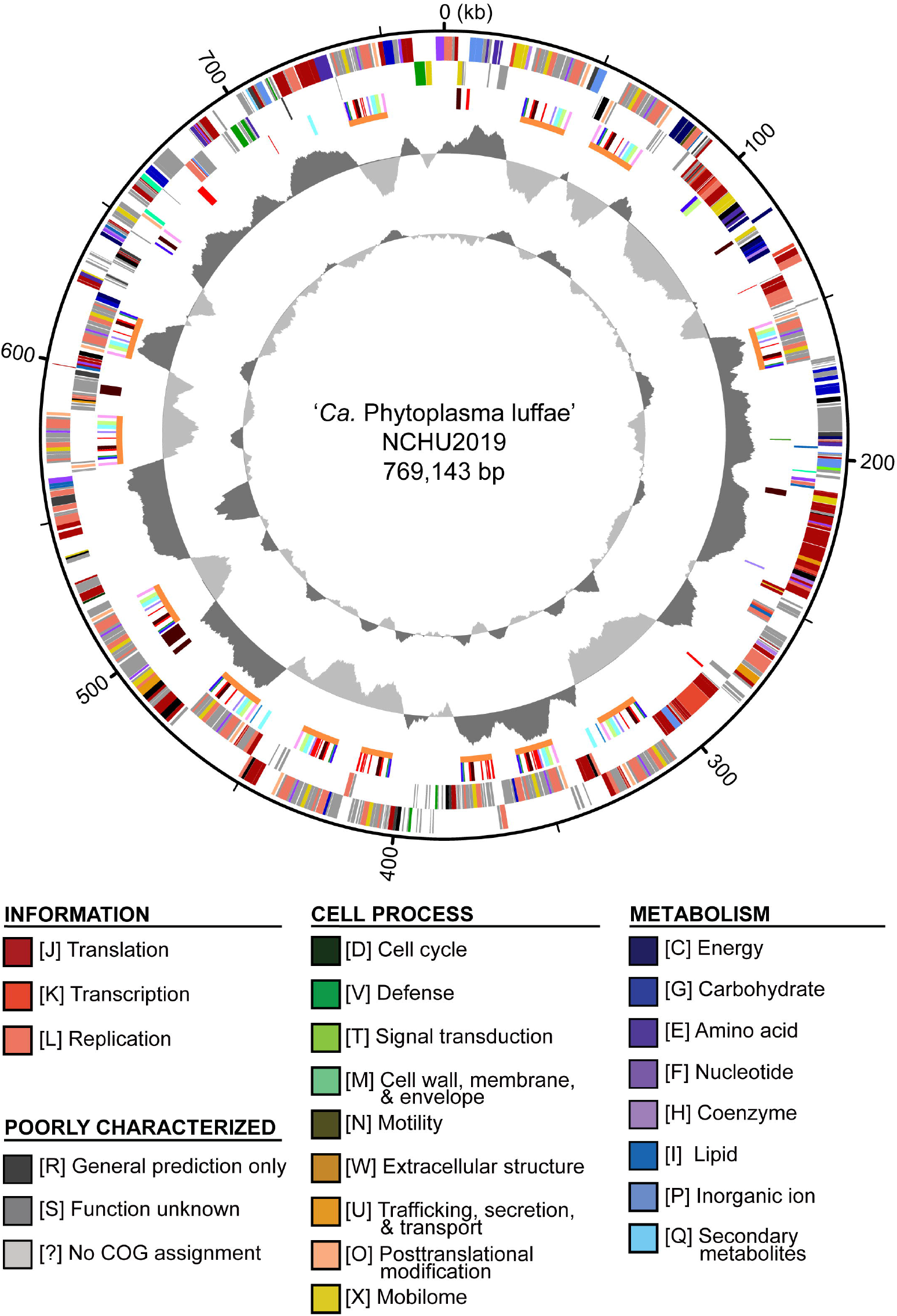
Genome map of ‘*Candidatus* Phytoplasma luffae’ NCHU2019. Rings from outside in: (1) Scale marks (kb). (2 and 3) Coding sequences on the forward and reverse strand, respectively. Color-coded by functional categories. (4) Genes associated with potential mobile units (PMUs), color-coded by annotation (see **Figure 5**). Gene clusters that represent individual PMUs are labeled by orange lines. Genes encode putative secreted proteins are in red. (5) GC skew (positive: dark gray; negative: light gray). (6) GC content (above average: dark gray; below average; light gray). One high GC peak located in the 531 to 544 kb region corresponds to two adjacent rRNA operons.

For genome size estimation based on k-mer analysis, we utilized k = 21 as a representative data set for visualization (**Figure S2A**). We found that while the peak depth is 51, the frequency distribution is nearly flat in the range between ~40-55. Based on this distribution, the genome size is estimated to be in the range between 660 and 842 kb (**Figure S2B**). The assembled genome size of 769 kb is near the middle point (i.e., 751 kb) of these estimates.

The annotation of this phytoplasma genome contains two complete sets of 16S-23S-5S rRNA gene clusters, 31 tRNA genes, 725 coding sequences (CDSs), and 13 pseudogenes. Both copies of the 16S rRNA gene are 100% identical to the reference sequence of ‘*Ca*. P. luffae’ LfWB (GenBank accession AF248956). Among the CDSs, 317 (44%) lacked any COG functional category assignments. Among the CDSs that were assigned to specific functional categories, those assigned to information storage and processing (e.g., replication, transcription, and translation) represent the largest group that account for 32% of all CDSs. In comparison, those assigned to cell process and metabolism account for 9% and 13%, respectively. These observations are consistent with findings from characterization of other phytoplasmas and more distantly related Mollicutes (e.g., *Spiroplasma*, *Entomoplasma*, and *Mycoplasma*) (Chen et al., 2012; Kube et al., 2012; Lo et al., 2018). The observation that a large fraction of genes lack specific functional annotation may be attributed to the elevated evolutionary rates of Mollicutes and their distant phylogenetic relationships from model organisms (Wu and Eisen, 2008; Wu et al., 2009; McCutcheon and Moran, 2012). The observation that genes related to information storage and processing genes are relatively abundant compared to those involved in metabolism is common among symbiotic bacteria with small genomes (McCutcheon and Moran, 2012; Lo et al., 2016).

Other than the low G+C content and reduced gene content, phytoplasma genomes are generally known to be repetitive, partly due to the presence of PMUs (Sugio and Hogenhout, 2012). Interestingly, the genome of this strain is far more repetitive than other phytoplasmas that have been characterized. On average, PMU core genes account for ~4.7% (Std. Dev. = 2.7%) of the genome size among those 19 representative phytoplasma strains analyzed (**Table 1**). For strain NCHU2019, there are 117 PMU core genes that are organized into at least 13 distinct PMU regions (**Figure 2**) and account for 11% of the chromosome length. Additionally, a pair of chromosomal segments, each is ~75 kb in size, were found to be duplicated (positions: 315,975-391,140 and 391,446-466,612). Together, these two duplicated segments and the 13 PMU regions account for 25% of the chromosome length. Explanation for the high genome repetitiveness of strain NCHU2019 compared to other phytoplasmas is unclear.

### Comparisons with Other Phytoplasmas

Based on the established classification scheme and a previous study of 16S rRNA gene phylogeny (Davis et al., 2017), ‘*Ca*. P. luffae’ belongs to group 16SrVIII and is most closely related to ‘*Ca*. P. malaysianum’ (group 16SrXXXII; GenBank accession EU371934) (Nejat et al., 2013). These two species-level taxa have 95.9% sequence identity (i.e., 1,463/1,526 aligned nucleotides) in their 16S rRNA genes. However, no genome sequence is available for ‘*Ca*. P. malaysianum’ for comparative analysis.

Other than ‘*Ca*. P. malaysianum’, the next closest relatives of ‘*Ca*. P. luffae’ include phytoplasmas belonging to groups 16SrV (‘*Ca*. P. vitis’ and ‘*Ca*. P. ziziphi’), 16SrVI (‘*Ca*. P. sudamericanum’ and ‘*Ca*. P. trifolii’), and 16SrVII (‘*Ca*. P. fraxini’). Among these, one complete genome sequence (GenBank accession CP025121) is available for ‘*Ca*. P. ziziphi’ strain Jwb-nky (Wang et al., 2018a), which represents the most closely related lineage for comparative analysis (**Figure 3A**). Comparison based on the 16S rRNA gene sequences indicated that ‘*Ca*. P. luffae’ NCHU2019 and ‘*Ca*. P. ziziphi’ Jwb-nky have 94.9% sequence identity (i.e., 1,447/1,524 aligned nucleotides). For whole-genome comparison, only 49% of the chromosomal segments can be mapped between these two strains and these segments have < 80% ANI. Pairwise genome alignment indicated that the most of the conserved regions correspond to PMUs (**Figure 4**), which further supports that the sequence divergence between these two genomes is too high for nucleotide-level comparisons. The lack of chromosome-level synteny conservation was commonly reported in previous comparisons between complete genome sequences of phytoplasmas (Bai et al., 2006; Tran-Nguyen et al., 2008; Andersen et al., 2013; Orlovskis et al., 2017; Wang et al., 2018a), even for ‘*Ca*. P. asteris’ strains sharing > 99.9% 16S rRNA gene sequence identity and > 98.1% genome-wide ANI (Cho et al., 2020a). These observations may be attributed to the strong nucleotide composition biases, the high mutation accumulation rates, and the influence of PMUs (Cho et al., 2020a).

**Figure 3.**
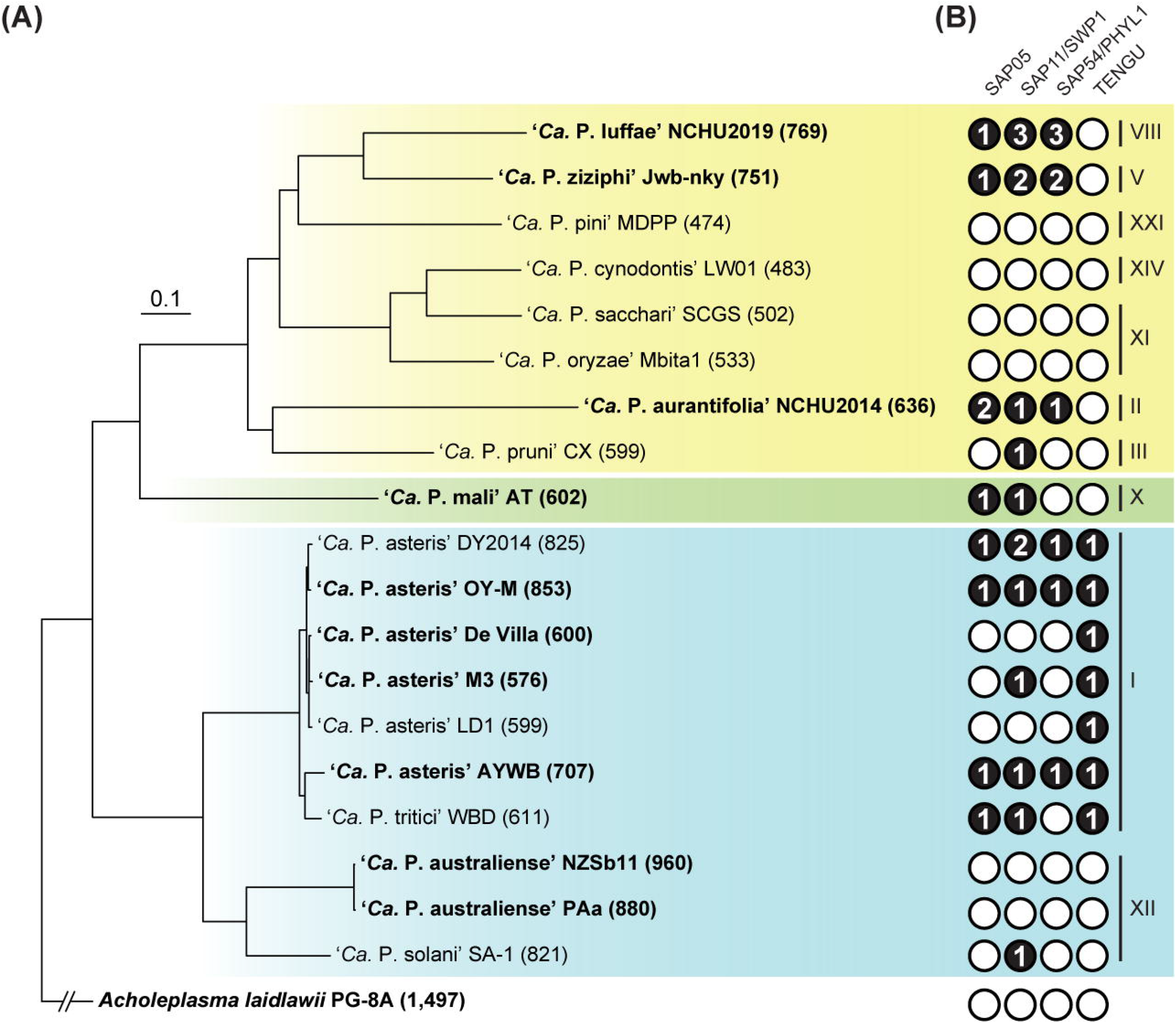
Evolutionary relationships among representative phytoplasmas with genome sequences available. The 16S rRNA gene (16Sr) group assignments are labeled on the right. The three major phylogenetic clusters are indicated by colored backgrounds (I: blue; II; green; III: yellow). *Acholeplasma laidlawii* is included as the outgroup. (A) Maximum likelihood phylogeny inferred based on a concatenated alignment of 80 conserved single-copy genes (27,742 aligned amino acid sites). All internal nodes received > 80% bootstrap support based on 1,000 re-sampling. Strains with complete genome assemblies available are highlighted in bold. The number in parentheses following the strain name indicates the genome size (in kb). (B) Distribution of known effector genes. Gene presence and absence are indicated by filled (with copy numbers) and empty circles, respectively.

**Figure 4.**
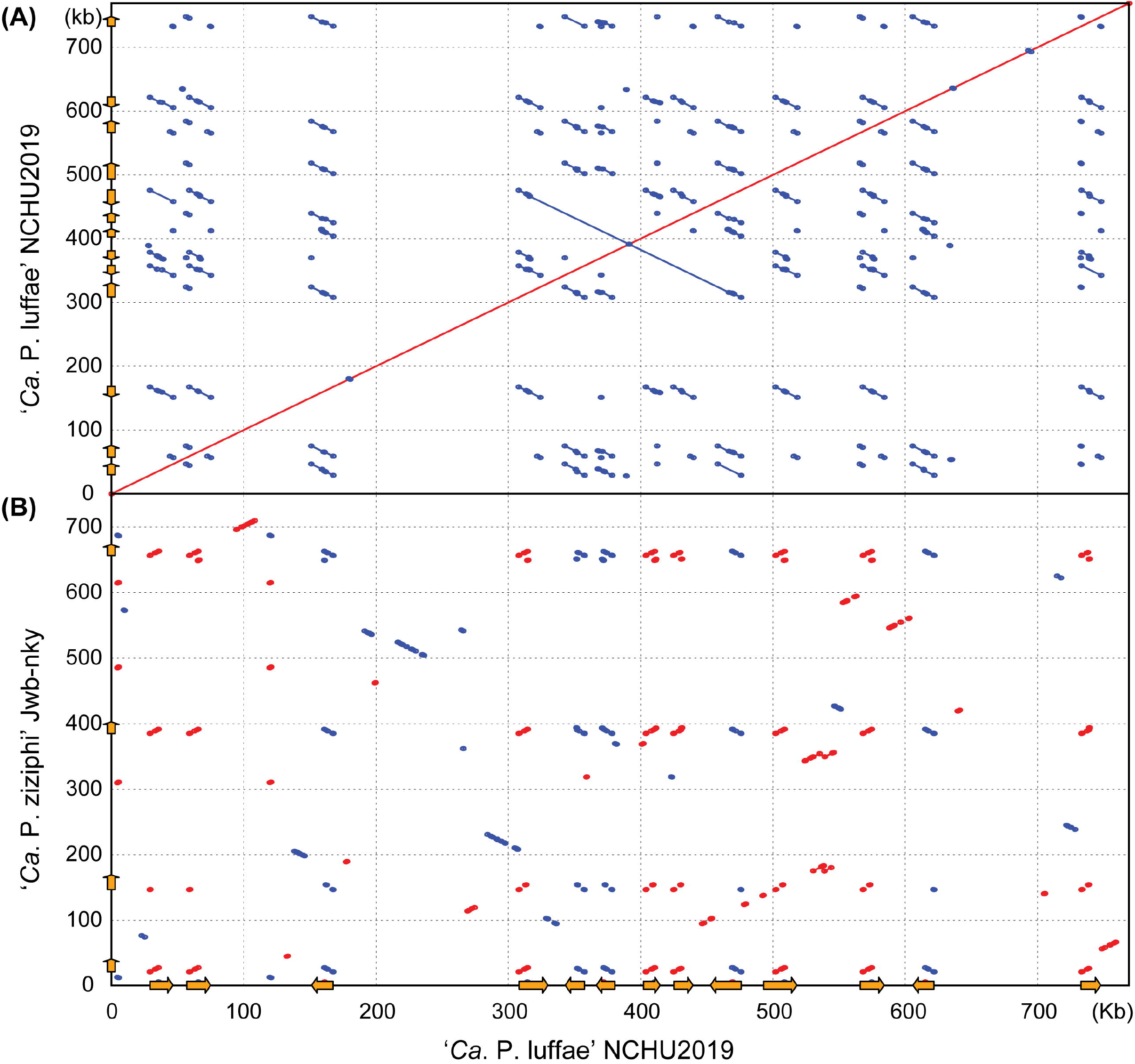
Pairwise genome alignments. The genome of ‘*Ca*. P. luffae’ NCHU2019 was used as the reference for comparison with (A) itself and (B) ‘*Ca*. P. ziziphi’ Jwb-nky. Matches on the same strand and the opposite strand are indicated in red and blue, respectively. Potential mobile units (PMUs) are illustrated as orange arrows on both axes.

Considering the high level of nucleotide sequence divergence between ‘*Ca*. P. luffae’ NCHU2019 and ‘*Ca*. P. ziziphi’ Jwb-nky, we also examined gene content based on protein sequence comparisons. In the pairwise comparison, 344 homologous gene clusters are shared between these two strains, while 175 and 241 are specific to NCHU2019 and Jwb-nky, respectively. These counts of shared and strain-specific genes are comparable to those found in previous studies of phytoplasma genome comparisons at between-species level (Chung et al., 2013; Tan et al., 2021). Although relatively large numbers of strain-specific genes were identified, > 80% of these genes lack functional annotation, making the inference of their roles difficult. As such, the gene content regarding metabolic capacity and transporters of ‘*Ca*. P. luffae’ NCHU2019 is expected to be highly similar to that of ‘*Ca*. P. ziziphi’ Jwb-nky, which was described in detail previously (Wang et al., 2018a).

For comparisons with those more divergent phytoplasmas with genomic information available (**Table 1**), only 134 homologous gene clusters are conserved among the 19 representatives analyzed. This estimate of phytoplasma core genome is much lower than the ^~^200 genes consistently reported in previous studies (Chen et al., 2012; Chung et al., 2013; Tan et al., 2021), an observation that is likely explained by the inclusion of several draft assemblies in this study. Detailed description regarding the functions of these ~200 core genes were reported previously (Chen et al., 2012; Kube et al., 2012; Chung et al., 2013).

### Detailed Characterization of PMUs

To better understand the roles of PMUs in phytoplasma genome evolution, we conducted detailed characterization of these mobile genetic elements. Among the 13 PMUs identified in ‘*Ca*. P. luffae’ NCHU2019 (**Figure 5A**), 11 are considered as complete ones and range from 14 to 18 kb in size. The remaining two (i.e., #6 and #7) are both 12 kb in size and appear to be truncated based in the lack of multiple PMU core genes (i.e., *tmk, dnaB, dnaG*,and *tra5*). These 13 PMUs are variable in the gene content in between *rad50* and *tmk*, while the sequences of shared core genes are highly conserved. For example, the *tmk* and *dnaB* homologs among these PMUs have identical sequences (**Figure 6**). It is unclear if the sequence conservation of these core genes is explained by lack of mutation accumulation, purifying selection, or assembly artifacts. Regarding chromosomal locations, these PMUs are interspersed across the entire chromosome and there is no obvious pattern of clustering (**Figures 2 and 4**). Two sets of PMUs (i.e., #4-6 and #7-9) are associated with the two 75-kb repeat regions. Notably, one set of junctions for these large repeats (chromosomal positions 315,975 and 466,612) are located inside PMU #4 (positions 308,032..323,983) and PMU #9 (positions 458,604..475,923). This finding suggests that homologous recombination facilitated by PMUs may have facilitated the segmental duplication of this chromosome, which in turn further increased the PMU copy number.

**Figure 5.**
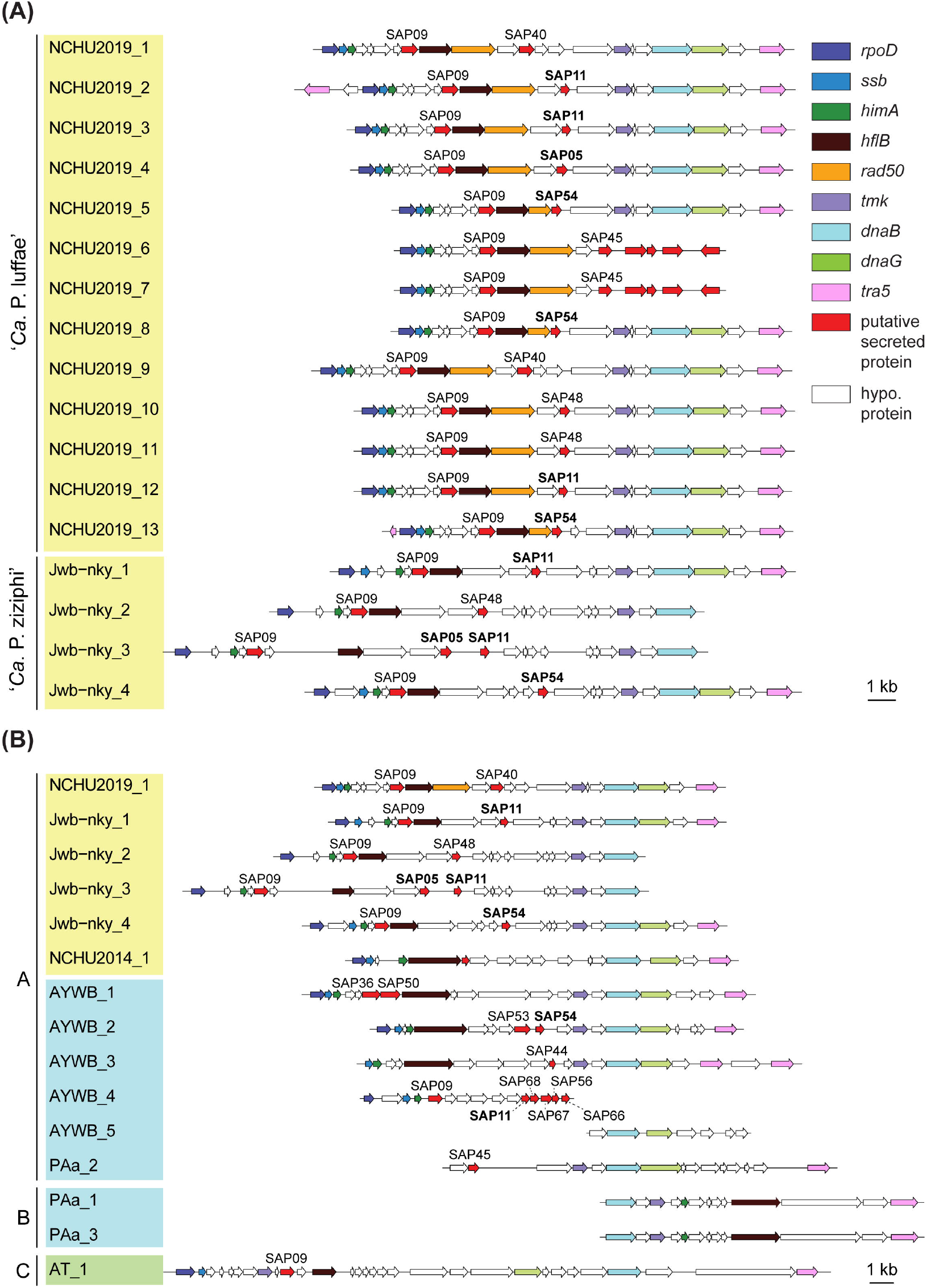
Gene organization of potential mobile units (PMUs). Each individual PMU is labeled by the phytoplasma strain name and a numerical identifier. Background colors for the PMU identifiers indicate the three phylogenetic clusters of phytoplasmas (I: blue; II; green; III: yellow). Genes are drawn to scale and color-coded according to annotation. Homologs of putative effectors identified in the ‘*Ca*. P. asteris’ AYWB genomes are labeled, those with experimental evidence (i.e., SAP05, SAP11, and SAP54) are highlighted in bold. (A) All PMUs in ‘*Ca*. P. luffae’ NCHU2019 and ‘*Ca*. P. ziziphi’ Jwb-nky. Grouped by genomes. (B) Representative PMUs from selected phytoplasmas. Grouped by PMU types.

**Figure 6.**
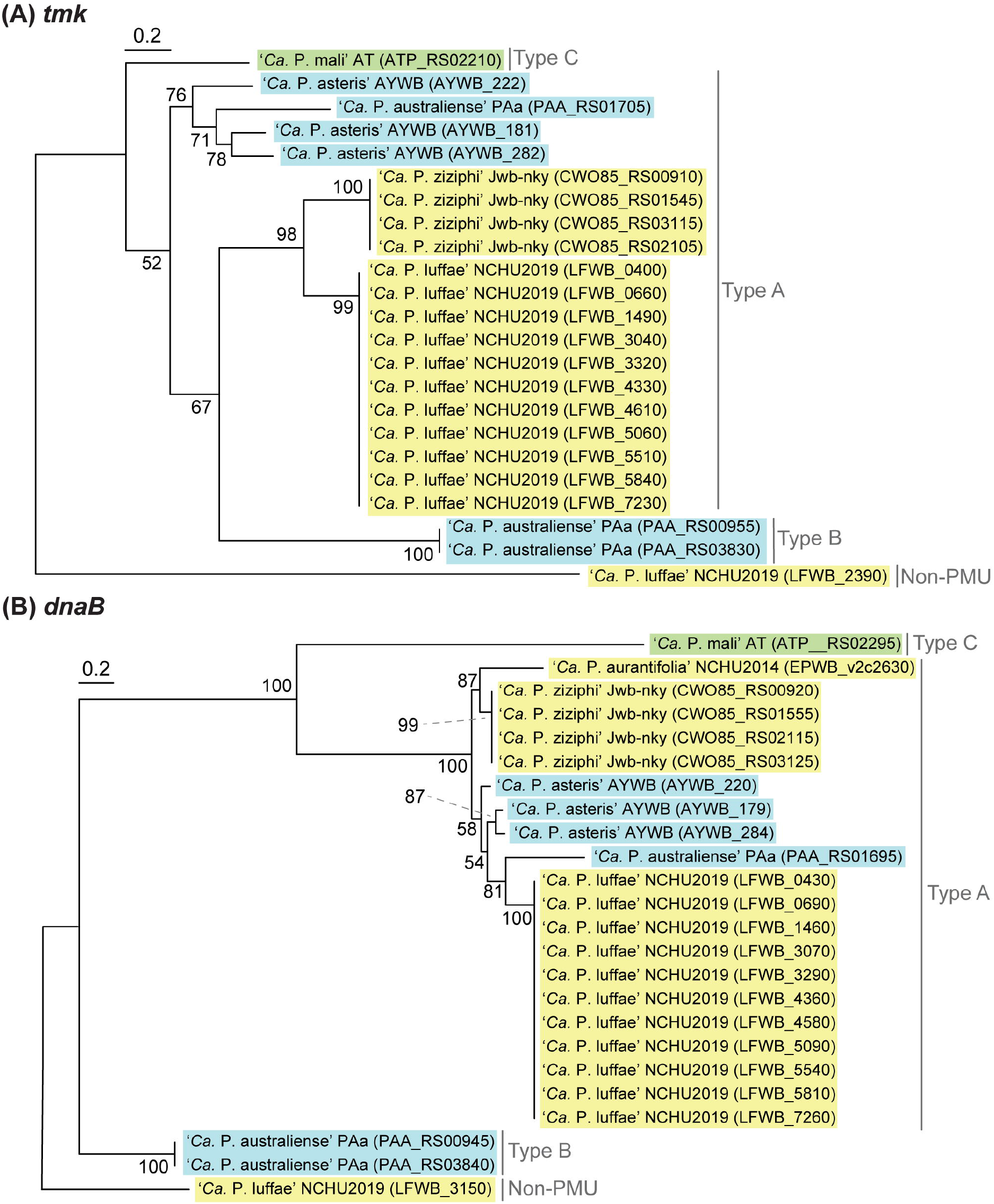
Maximum likelihood phylogenies of PMU core genes (A) *tmk* (228 aligned amino acid sites). (B) *dnaB* (531 aligned amino acid sites). In both panels, a non-PMU homolog is included as the outgroup. Numbers next to internal branches indicate the bootstrap support levels based on 1,000 re-sampling. Background colors for the gene identifiers indicate the three phylogenetic clusters of phytoplasmas (I: blue; II; green; III: yellow).

Based on initial characterization of the PMUs in ‘*Ca*. P. asteris’ AYWB genome, eight core genes were identified (Bai et al., 2006). As more genome sequences become available from more diverse phytoplasmas, we are able to include representatives of 13 ‘*Ca*. P.’ species from all three phylogenetic clusters in this analysis (**Table 1**). Compared to other phytoplasmas, ‘*Ca*. P. luffae’ NCHU2019 has the highest number of PMUs and is distinctive in that all of its PMUs are similar (**Figure 5A**). Among the 11 complete PMUs in this genome, the eight PMU core genes and *rad50* are all organized in the same order. The minor variation in gene organization is mainly in between *rad50* and *tmk*, where genes encode different putative secreted proteins and hypothetical proteins are found. For comparison, in the closely related ‘*Ca*. P. ziziphi’ Jwb-nky of cluster III (i.e., the yellow clade in **Figure 3A**), the four PMUs exhibit much higher levels of diversity in gene organization between *hflB* and *tmk* (**Figure 5A**). For the distantly related ‘*Ca*. P. asteris’ AYWB and ‘*Ca*. P. australiense’ PAa of cluster I (i.e., the blue clade in **Figure 3A**), high levels of intra-genomic PMU diversity are also observed (**Figure 5B**).

Based on the presence/absence pattern and order of eight PMU core genes defined previously (Bai et al., 2006), we found that the PMUs in these representative phytoplasmas with complete genome sequences available can be classified into three major types (**Figure 5B**). The strains omitted in the visualization all have close relatives belonging to the same 16Sr group and share similar PMUs. Among the three major types, type A PMUs, in which *tmk* is upstream of *dnaB*, are the most common ones that include PMUs found in phylogenetic clusters I (e.g., ‘*Ca*. P. asteris’ and ‘*Ca*. P. australiense’) and III (e.g., ‘*Ca*. P. luffae’, ‘*Ca*. P. ziziphi’, and ‘*Ca*. P. aurantifolia’). Type B PMUs are shorter, have *tmk* downstream of *dnaB*, and are found only in ‘*Ca*. P. australiense’ that belongs to cluster I. Type C is the rarest type, with only one representative found in ‘*Ca*. P. mali’ that belongs to cluster II, and has *hflB* and *dnaG* located in between *tmk* and *dnaB*. In addition to the gene organization, molecular phylogenies of *dnaB* and *tmk* also revealed divergence among homologs from different PMU types (**Figure 6**). These patterns provide further support for our PMU classification scheme. It is interesting to note that regardless of PMU copy numbers, most of the phytoplasmas with genome sequences available harbor only one single type of PMUs. ‘*Ca*. P. australiense’ is the only exception that harbors both type A and B PMUs. Future improvements in sampling more diverse phytoplasma genomes, particularly cluster II lineages, are necessary to provide a more comprehensive understanding of PMU diversity.

### PMU and Phytoplasma Genome Size Variation

One notable observation regarding phytoplasma genomes is the extensive size variation at both between- and within-species levels (**Table 1**). Phylogenetic relatedness, as quantified by core genome sequence divergence, does not appear to provide a reliable predictor for genome sizes (**Figure 3A**). Previous within-species comparisons suggested that PMU abundance is an important factor (Bai et al., 2006; Andersen et al., 2013). To further test if this pattern holds true for genus-level analysis, we performed regression analysis to examine the correlation between the combined length of PMU core genes and genome size. Strikingly, when all 19 representative phytoplasma genome assemblies were examined together, the combined length of PMU core genes explains 79% of the variance in genome sizes (*r* = 0.89, *p* = 3.7e-07) (**Figure 7**). Due to the concern that draft assemblies cannot provide accurate information regarding these two metrics, we also performed regression analysis with only the 10 complete assemblies and obtained similar results (*r* = 0.87, *p* = 9.7e-04).

**Figure 7.**
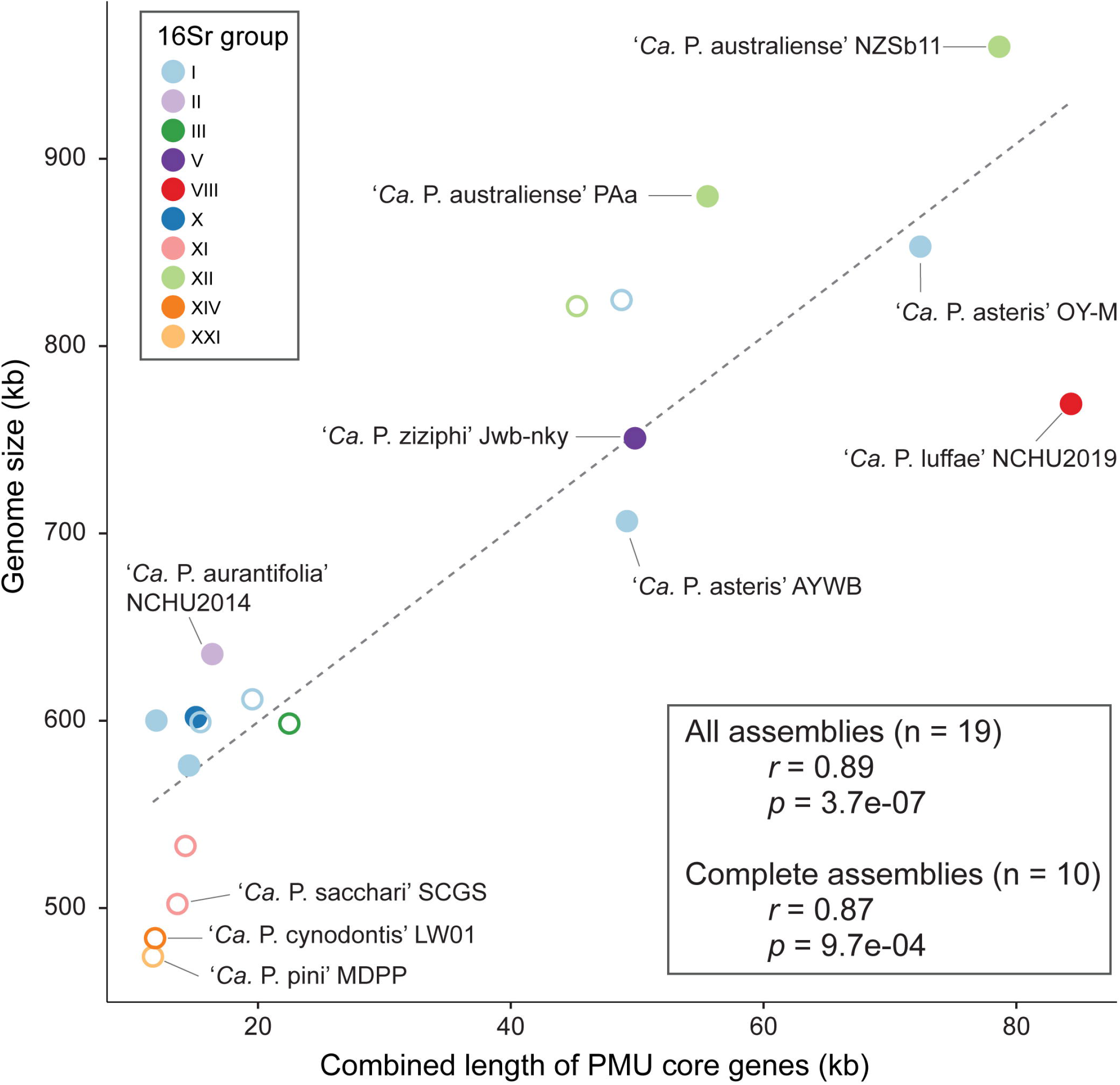
Correlation between the combined length of PMU core genes and genome size. Strains with complete and draft genome assemblies are indicated by filled and open circles, respectively. The linear regression line was based on all available assemblies.

The evolution of bacterial genome sizes is a topic that received much research attention and previous studies have identified multiple relevant factors, such as composition of gene content, prevalence of mobile genetic elements, effective population sizes, level of mutational biases toward deletion, and ecological niches (Mira et al., 2001; Konstantinidis and Tiedje, 2004; Ochman and Davalos, 2006; Kuo and Ochman, 2009; Kuo et al., 2009; Novichkov et al., 2009; McCutcheon and Moran, 2012; Lo et al., 2016; Sabater-Muñoz et al., 2017; Weinert and Welch, 2017). Due to the complexity of bacterial genome size evolution and the multitude of factors involved in the process, one single factor (i.e., PMU abundance) can have such strong correlation with phytoplasma genome size is surprising. Notably, as genome reduction appears to be a common and recurring theme of symbiont evolution, the roles of PMU in genome expansion of some phytoplasmas require further investigation. Because PMUs are known to be associated with effector genes and likely can transfer horizontally between closely- (Cho et al., 2019; Seruga Music et al., 2019) or distantly-related phytoplasmas (Chung et al., 2013; Ku et al., 2013), the involvement of PMUs in phytoplasma effector gene content evolution is particularly important. However, it is worth noting that even though horizontal transfers of PMUs may provide novel combination of effector gene content, PMU evolution may have been dominated by vertical inheritance or at least transfers between closely related lineages based on the observation that in most cases related lineages have similar PMUs.

### Effector Genes

An important feature of phytoplasmas is their ability to modulate host plant development through effectors, which are small secreted proteins (Sugio et al., 2011b; Sugio and Hogenhout, 2012). To date, four phytoplasma effectors have been experimentally characterized, including SAP05 (Gamboa et al., 2019; Huang and Hogenhout, 2019; Huang et al., 2021), SAP11/SWP1 (Bai et al., 2009; Sugio et al., 2011a; Lu et al., 2014; Chang et al., 2018; Wang et al., 2018c, 2018b), SAP54/PHYL1 (MacLean et al., 2011; Maejima et al., 2014; Orlovskis and Hogenhout, 2016), and TENGU (Hoshi et al., 2009; Sugawara et al., 2013; Minato et al., 2014). The expanded genome sequence availability allowed us to investigate the phylogenetic distribution of homologous effector genes among diverse phytoplasmas. The highly variable distribution patterns suggest that the effector gene content may have rapid turnover during the diversification of phytoplasmas (**Figure 3**). For example, at within-species level comparison among ‘*Ca*. P. asteris’ strains, the patterns of presence/absence and gene copy number are variable for three of these effectors (**Figure 3B**) although the level of core genome sequence divergence is very low as indicated by the short branch lengths (**Figure 3A**). This finding is consistent with our previous in-depth characterization of 16SrI phytoplasmas (Cho et al., 2020a). At genus-level, TENGU appears to be restricted to and conserved among 16SrI lineages in cluster I (i.e., ‘*Ca*. P. asteris’ and ‘*Ca*. P. tritici’), while the other three effectors are variable. Based on these patterns, it is likely that TENGU originated in the most recent common ancestor (MRCA) of 16SrI phytoplasmas. However, for the other three effectors, it is unclear if the MRCA of all extant phytoplasmas harbor these genes or not. If yes, then multiple independent gene losses are required to explain the distribution of these genes among extant phytoplasmas. Alternatively, multiple independent origins, likely mediated by PMU-mediated horizontal gene transfer (Chung et al., 2013; Cho et al., 2019; Seruga Music et al., 2019), are required to explain the gene distribution pattern.

Another interesting observation is that several phytoplasmas do not possess any of these four effector genes. It is likely that these phytoplasmas harbor novel effector genes that are yet to be characterized and further investigation is required to obtain a more complete picture of phytoplasma effector diversity.

For closer inspection of these four known effectors, we performed multiple sequence alignments to examine the protein sequence divergence among homologs (**Figure S3**). Consistent with the expectation derived from their phylogenetic distribution patterns, the three effector genes found among diverse phytoplasmas have higher levels of sequence divergence compared to phylogenetically restricted TENGU homologs.

For copy number variation, ‘*Ca*. P. luffae’ NCHU2019 stands out as having the highest copy numbers for SAP11 and SAP54 (**Figure 3B**). These homologous genes are all located within PMUs (**Figure 5**) and have nearly identical sequences (**Figure S3**), which suggest that recent intra-genomic PMU duplications are responsible for expansions in effector gene copy numbers. Similar patterns were observed for SAP11 and SAP54 homologs in ‘*Ca*. P. ziziphi’ Jwb-nky. Experimental characterization of the two SAP11 homologs in ‘*Ca*. P. ziziphi’ demonstrated that both can stimulate lateral bud outgrowth for witches’ broom symptoms when expressed in *Nicotiana benthamiana* (Zhou et al., 2021). Intriguingly, in phytoplasma-infected jujube plants, these two SAP11 homologs have different expression patterns among tissue types. These findings suggest that such gene duplication events may lead to neofunctionalization or subfunctionalization, thus promoting the genetic diversity of effectors.

## Conclusion

In this work, we determined the complete genome sequence of an uncultivated ‘*Ca*. P. luffae’ strain associated with the witches’ broom disease of loofah. This assembly provides the first representative genome sequence for the 16SrVIII group of phytoplasmas and improves the taxon sampling of these diverse plant-pathogenic bacteria. For comparative genomics analysis conducted at genus level, we provided a global view of the PMUs (i.e., phytoplasma-specific mobile genetic elements) and identified three major PMU types that differ in gene organization and phylogenetic distribution. Importantly, statistical analysis revealed that PMU abundance explains nearly 80% of the variance in phytoplasma genome sizes, thus providing a quantitative estimate on the importance of these elements. Finally, our investigation of effector genes highlighted the genetic diversity associated with phytoplasma virulence and established the roles of PMUs in shaping such diversity.

## Data Availability

This genome sequencing project was deposited in the NCBI under BioProject PRJNA636624. The raw reads were deposited in the Sequence Read Archive (SRA) under the accessions SRR11921288 (Illumina MiSeq paired-end reads) and SRR11921289 (ONT MinION reads). The complete and annotated genome sequence of ‘*Ca*. P. luffae’ NCHU2019 was deposited in GenBank under the accession CP054393.

## Author Contribution

JYY and CHK conceptualized the study, acquired funding, and supervised the project. JYY, YCC, and CMT provided the biological materials. CMT and STC coordinated the sequencing. CMT and STC conducted the initial genome assembly. STC completed the assembly. YCL and CTH validate the assembly. STC and CTH conducted the comparative analysis and prepared the figures. CTH and CHK wrote the draft. All authors approved the submitted version.

## Funding

Work in the Yang lab was supported by grants-in-aid from the Ministry of Science and Technology (110-2628-B-005-002) and the Advanced Plant Biotechnology Center from the Featured Areas Research Center Program within the framework of the Higher Education Sprout Project by the Ministry of Education in Taiwan. Work in the Kuo lab was supported by Academia Sinica and the Ministry of Science and Technology (106-2311-B-001-028-MY3) of Taiwan. The funders had no role in study design, data collection and interpretation, or the decision to submit the work for publication.

## Acknowledgments

We thank Ai-Ping Chen and Shu-Jen Chou for technical assistance. The Illumina sequencing library preparation service was provided by the Genomic Technology Core Facility (Institute of Plant and Microbial Biology, Academia Sinica, Taipei. Taiwan). The Illumina paired-end sequencing service was provided by the Genomics Core Facility (Institute of Molecular Biology, Academia Sinica, Taipei. Taiwan).

## Conflict of Interest Statement

The authors declare that the research was conducted in the absence of any commercial or financial relationships that could be construed as a potential conflict of interest.

## Supplementary Materials

**Figure S1.**Coverage pattern of raw read mapping results to the genome assembly. Rings from outside in: (1) Scale marks (kb). (2) Sequencing depth calculated from Illumina (dark blue; outward) and ONT (light blue; inward) raw reads mapped to the assembly using a sliding window of 200-bp. An alignment score cutoff of 100 was applied to remove spurious hits. The peaks located in the 531 to 544 kb region correspond to rRNA gene clusters. (3) Potential mobile unit (PMU) regions (orange). (4) Two duplicated chromosomal segments (green; positions: 316-391 and 391-467 kb).

**Figure S2.**Genome size estimation based on k-mer analysis. (A) A representative frequency distribution of k-mer depth (k = 21). The peak depth is 51. (B) Correspondence between genome size estimates and k-mer depths.

**Figure S3.**Protein sequence alignments of four characterized phytoplasma effectors.

Individual effectors are identified using locus tags. Shaded background colors indicate the levels of sequence conservation. (A) SAP05. (B) SAP11/SWP1. (C) SAP54/PHYL1. (D) TENGU.

**Data S1.**Concatenated multiple sequence alignment of conserved single copy-genes used for inferring the molecular phylogeny shown in Figure 3. Simple text file in PHYLIP format.

## Notes

### Competing Interest Statement

The authors have declared no competing interest.

