## Supplementary figures and images for "Comparative Genome Analysis of ‘*Candidatus* Phytoplasma luffae’ Reveals the Influential Roles of Potential Mobile Units in Phytoplasma Evolution"

### Figure S1

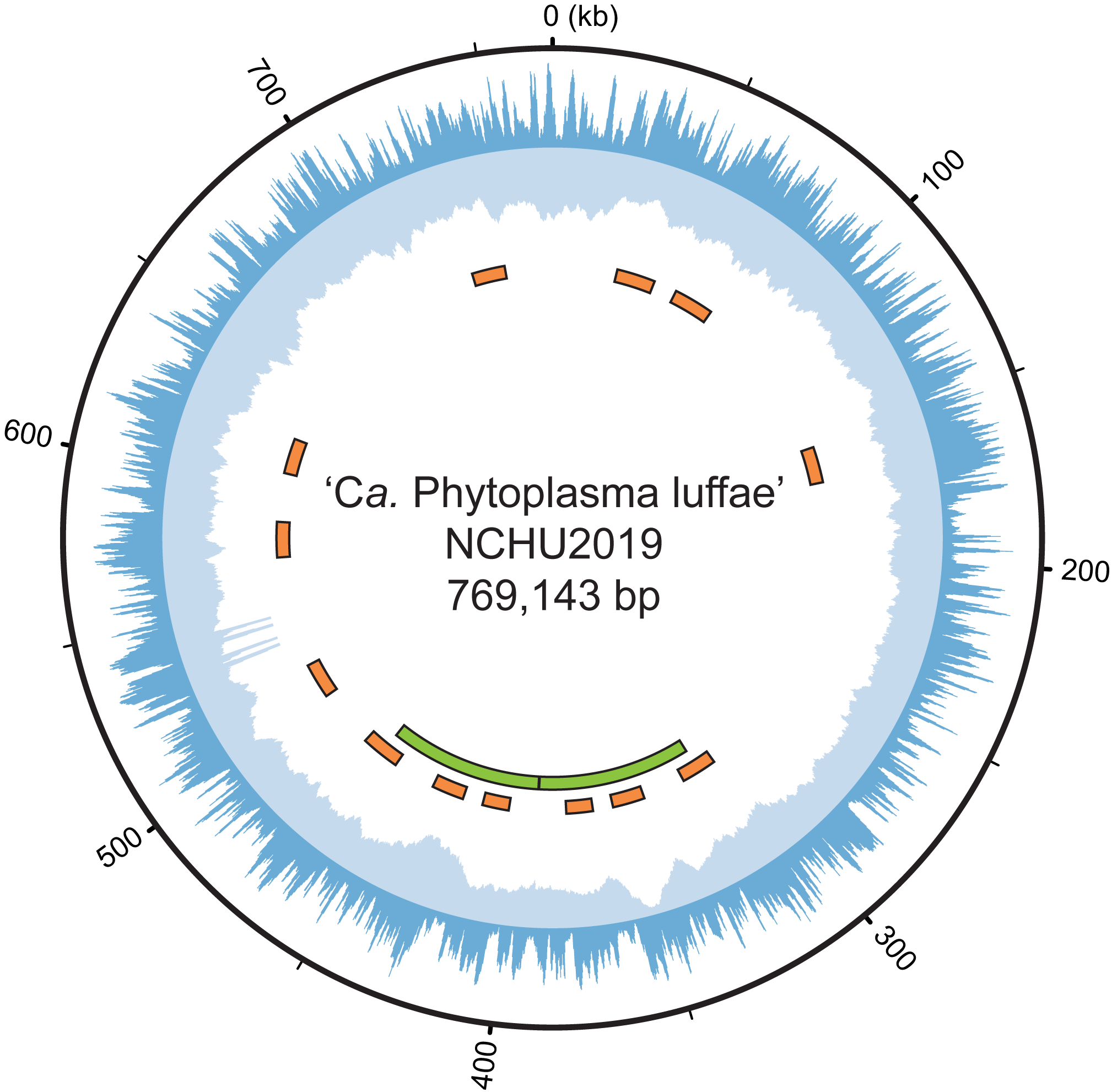

### Figure S2

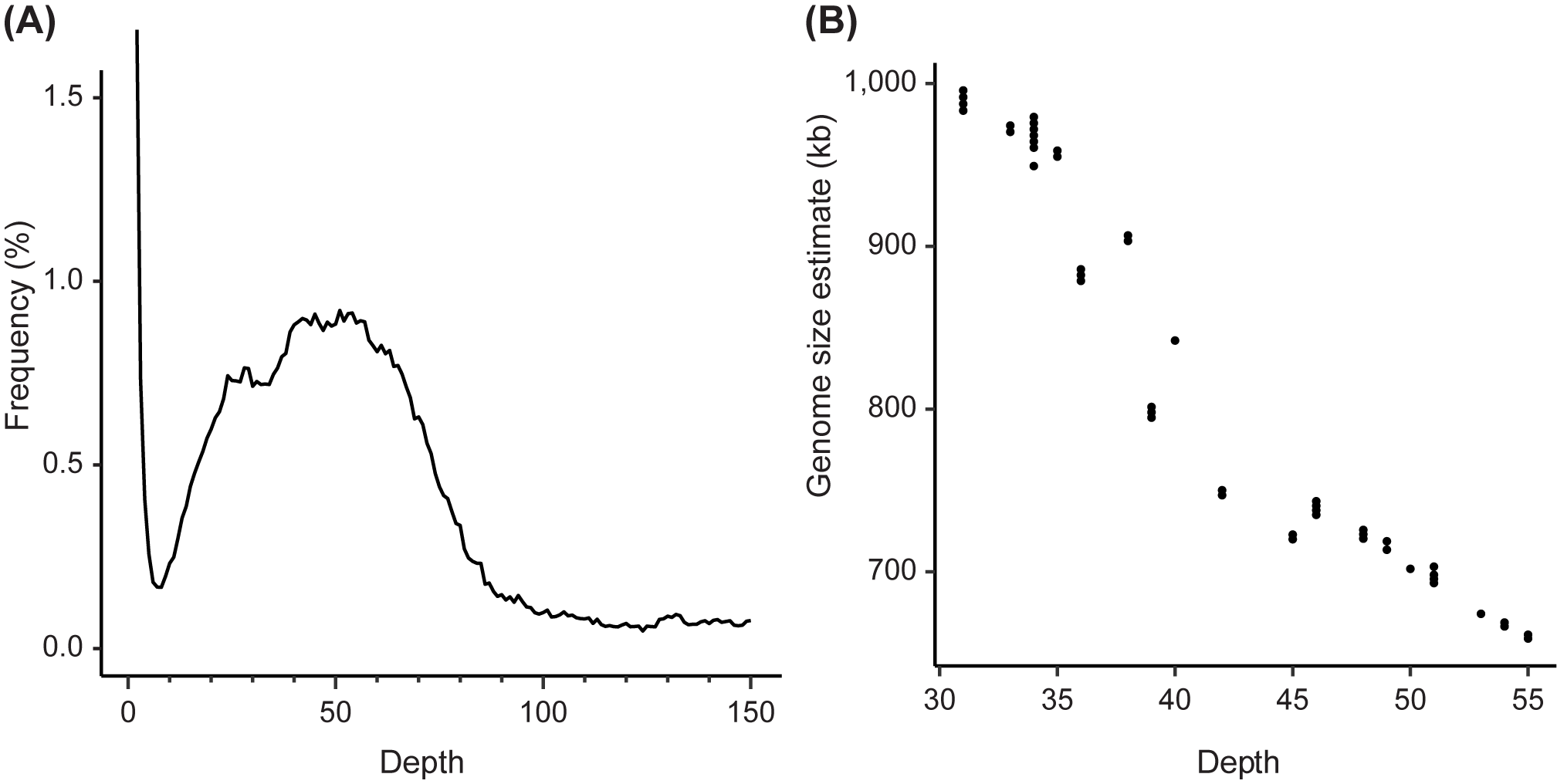

### Figure S3

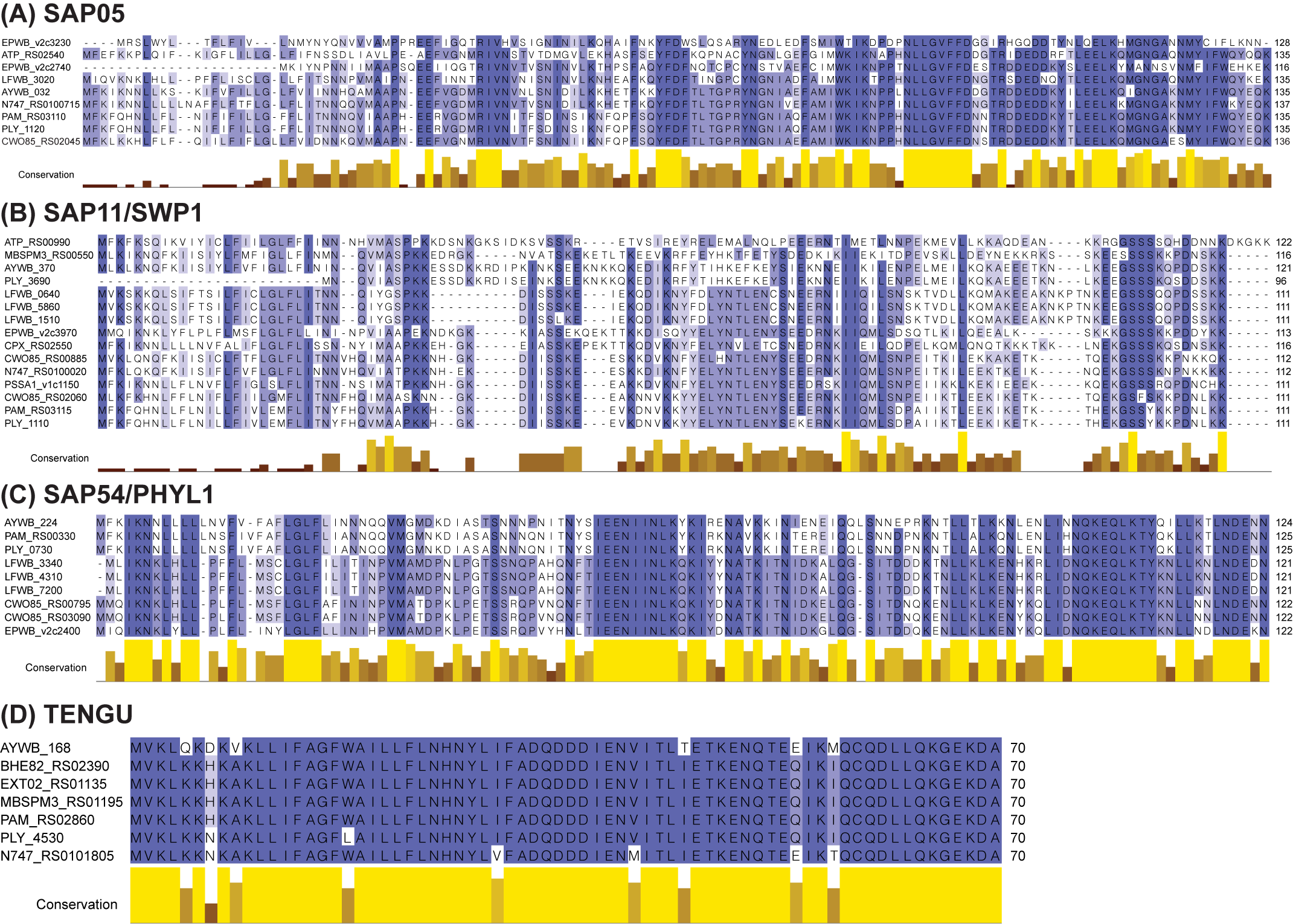
